# PTMExplorer: A Multi-Dimensional Integrative Visualization Platform for Protein Post-Translational Modification Function and Structure

**DOI:** 10.64898/2026.09.21.753344

**Authors:** Chenxia Li, Jinhao Wang, Chongyang He, Huqiang Wang, Sisi Geng, Yuanxiang Lao, Yupeng Zhang, Yunping Zhu, Songfeng Wu

## Abstract

Deciphering the functions of post-translational modifications (PTMs) is a critical bridge connecting large-scale modification proteomics data to mechanistic studies. However, most existing tools for visualizing PTM omics data are limited to site catalogs or single-dimensional feature displays. They lack the capability to simultaneously map user-derived differential modification sites onto multi-dimensional contexts, including protein three-dimensional (3D) structure, evolutionary conservation, functional sites, and disease associations. This limitation makes it difficult for researchers to rapidly assess the biological importance of candidate sites from among a vast number of differentially modified sites. Here, we present PTMExplorer, an interactive platform for the multi-dimensional visualization of protein PTMs. PTMExplorer comprises three core modules: PTM Inspector, built upon ProtVista, provides a multi-track, sequence-feature integrated view incorporating intrinsically disordered region (IDR) prediction (via flDPnn), surface accessibility calculation (via FreeSASA), and UniProt functional annotations; PTM 3D Locator, leveraging the Nightingale/Mol* engine, anchors modification sites onto AlphaFold/Protein Data Bank (PDB) 3D structures through residue mapping via PDBe-SIFTS; and PTM Overview, utilizing the R circlize package, presents a panoramic polar circos plot illustrating modification distribution and inter-group differential regulation. Additionally, three major disease-associated modification databases—PTMD, qPTM, and PhosCancer—are integrated as PTM-Disease Nexus, enabling co-localization comparison between user-defined differential sites and reported disease-related sites. PTMExplorer currently supports eight model organisms, accepts user-uploaded differential analysis results, and provides multi-dimensional annotations and various visualization options (https://www.bioladder.cn/PTMExplorer/). Using a multi-omics dataset from hepatocellular carcinoma (18 patients, 9 modification types) as a case study, we demonstrate the practical utility of PTMExplorer in screening potential biomarkers, revealing multi-modification coordination mechanisms, and distinguishing between absolute and relative quantification patterns.

## 1. Introduction

Post-translational modifications (PTMs) are fundamental to life, regulating protein function, stability, subcellular localization, and interaction networks through covalent addition or enzymatic cleavage^1^. Modifications such as phosphorylation, acetylation, ubiquitination, and glycosylation are extensively involved in critical biological processes including signal transduction, metabolic regulation, the cell cycle, and stress responses^2^. Aberrant PTMs are closely linked to the pathogenesis of major diseases like Alzheimer’s disease, Parkinson’s disease, and cancer^3,4^. For instance, over 55 phosphorylation sites have been identified on the tau protein, and dysregulation of its ubiquitination, acetylation, and methylation contributes to abnormal aggregation and neurofibrillary tangle formation, a hallmark pathology of Alzheimer’s disease^4^. Therefore, systematic characterization of PTM sites, types, and functional contexts is crucial for understanding disease mechanisms and discovering biomarkers.

Recent rapid advancements in high-resolution mass spectrometry and data acquisition strategies have propelled deep profiling in PTM proteomics. Four-dimensional (4D) proteomics, which adds ion mobility to retention time, mass-to-charge ratio, and intensity, significantly improves the sensitivity and accuracy of peptide identification^5^. Data-independent acquisition (DIA) workflows combined with deep learning prediction frameworks enable large-scale PTM site quantification without experimental spectral libraries^5^. The maturation of techniques like 4D-Label Free and DIA-PASEF allows researchers to systematically delineate differential landscapes of various modification types (e.g., phosphorylation, acetylation, succinylation) in clinical samples^6^. However, large-scale site identification is merely the starting point. Confronted with hundreds to thousands of differentially modified sites, the core bottleneck limiting research efficiency is how to rapidly prioritize biologically significant sites worthy of downstream functional validation.

The biological function of a PTM is highly dependent on its location within the protein’s structural and functional context. On the same protein, different modification sites may reside in intrinsically disordered regions (IDRs), active sites, protein-protein interaction interfaces, or surface-exposed areas, each carrying vastly different functional implications^7,8^. Sites located in active pockets or conserved regions often serve as critical nodes in signaling regulation, whereas those in disordered regions might represent stochastic events or noise^7^. Furthermore, the functional significance of a PTM should be evaluated in conjunction with disease context. Differential regulation of the same modification site between normal and tumor tissues is a key criterion for assessing its potential as a biomarker or therapeutic target^9^. Consequently, integrating PTM identification and differential analysis results with multi-dimensional features—such as domain architecture, conservation, surface accessibility, disorder propensity, functional sites, and disease associations—is essential for advancing from “data generation” to “mechanistic insight” in PTM functional studies.

To meet the needs of PTM data interpretation, numerous tools and databases have been developed. Based on their functionality, they can be broadly categorized into four classes.

### Catalog-type databases

represented by PhosphoSitePlus (https://www.phosphosite.org) and Phospho.ELM (http://phospho.elm.eu.org), primarily offer site directories. PhosphoSitePlus curates over 450,000 non-redundant modification sites, supporting filtering by disease, tissue, cell line, modification type, and domain^10^. Phospho.ELM contains over 42,000 phosphorylation sites, integrating disorder, solvent accessibility, and conservation scores, and provides kinase-substrate network links^11^. dbPTM 2025 includes over 2.79 million PTM sites from 48 data sources, covering proteomic data for 13 cancer types^12^. PhosCancer catalogues 174,587 phosphorylation sites across 12 cancer types, offering multi-dimensional annotations including 3D structure, functional domains, upstream kinases, and clinical features^13^. qPTM integrates experimentally validated PTM sites in the human proteome with cancer multi-omics data^14^. PTMD specializes in collecting disease-associated PTM sites and their regulatory mechanisms^15^. The primary limitation of these tools is their nature as catalog browsers; they cannot import user-specific differential analysis results to display modification abundance changes under specific experimental conditions (e.g., disease versus control).

### Analytical visualization tools

allow users to upload custom data for PTM analysis. Scop3P integrates UniProtKB sequences and PDB structures, mapping phosphorylation sites onto 1D sequence contexts and 3D structures, and provides circos plots showing biophysical features like early folding propensity and disorder probability^16^. However, Scop3P only supports phosphorylation. AlphaMap offers a Python package for mapping MS-identified peptides and PTMs onto protein sequences, overlaying UniProt annotation tracks, and supports input formats from mainstream search engines like MaxQuant, DIA-NN, and Spectronaut^17^. ProtVista, an EBI-developed JavaScript component for general protein feature visualization used by UniProt and the Open Targets Platform, is built on D3.js and supports multi-track feature display and click-highlighting within the sequence context^18^. Nevertheless, ProtVista itself only displays UniProt annotations and cannot directly import user-provided quantitative data. PTMVision, published in 2025, supports sample-level and protein-level visualization for various PTM types identified via open searches, integrating 3Dmol.js for 3D rendering and contact maps^19^. However, its 3D representation focuses on contact maps, lacks systematic integration with UniProt functional annotations, and does not support user-defined thresholds for differential analysis (PMID: 39772617).

### Structure-function integrative tools

have begun to associate PTMs with 3D protein structures. Missense3D-PTMdb, published in 2025, visualizes both missense variants and PTM sites on AlphaFold models of the human proteome, but its focus is on variant-structure mapping rather than PTM differential analysis^20^. MutationExplorer maps sequence mutations onto 3D structures and displays real-time Rosetta energy predictions, but targets amino acid substitutions, not PTMs^21^.

### Pathway and network tools

include PTMint and PhosNetVis. PTMint integrates experimentally validated PTM regulation of protein-protein interaction (PPI) networks^22^. PTMNavigator, published in 2025, projects differentially regulated PTMs onto KEGG/WikiPathways maps for pathway-level enrichment analysis^23^. PhosNetVis provides kinase enrichment analysis and 2D/3D network visualization^24^. None of these tools, however, provide an integrated view that simultaneously maps user-derived differential PTM sites onto 3D structures overlaid with functional annotations.

In summary, existing PTM visualization tools generally suffer from four key limitations that hinder comprehensive functional assessment:

1. **Lack of joint display for multiple modification types and protein functional features:** Most tools support only a single modification type (e.g., phosphorylation) or display a limited set of UniProt feature tracks, failing to integrate multi-dimensional protein characteristics like disorder, surface accessibility, conservation, and functional sites in a single view.
2. **Inability to localize PTMs at the 3D structural level:** Catalog tools and most analytical visualization tools do not offer spatial localization of modification sites on 3D structures. The few tools with 3D capabilities (e.g., PTMVision) do not systematically integrate UniProt domain and functional site annotations.
3. **Inability to import user differential analysis results:** Catalog databases fundamentally do not support uploading and displaying user-customized experimental data. Existing analytical tools (e.g., AlphaMap) mainly display MS identification information, lacking systematic visualization schemes for differential PTMs derived from disease comparisons or treatment groups.
4. **Lack of an integrated perspective for disease-modification co-localization:** Disease-related modification databases like PTMD, qPTM, and PhosCancer operate independently. Users cannot co-localize their own experimental differential PTMs with reported disease-associated modification sites within a single tool.

To address these challenges, we developed PTMExplorer—an interactive visualization platform that synchronously maps proteomic PTM identification and differential data onto multi-dimensional contexts, including protein domains, functional sites, 3D structures, and disease-modification associations. PTMExplorer is integrated into the BioLader bioinformatics cloud platform (https://www.bioladder.cn), which also facilitates data pre-processing for search engine results^25^.

PTMExplorer consists of three core interactive visualization modules:

- **PTM Inspector (Feature Landscape):** Built on the ProtVista component^18^, this module maps PTM sites onto a protein sequence track, overlaying multi-dimensional feature tracks such as UniProt-annotated domains and sites (e.g., active sites, binding sites), known PTM sites, predicted IDRs (using flDPnn, IDR score)^26^, and surface accessibility (RSA values calculated from AlphaFold structures via FreeSASA). This enables users to comprehensively assess the functional importance of modification sites within a one-dimensional sequence view.
- **PTM 3D Locator (Structure View):** Leveraging the Nightingale nightingale-structure component^27^ with the Mol* 3D rendering engine, this module anchors PTM sites onto corresponding AlphaFold or PDB 3D structures via residue-level mapping provided by PDBe-SIFTS. It visually highlights the spatial distribution of modification sites within the protein’s 3D space, aiding in determining whether a site is surface-exposed or located in a pocket region.
- **PTM Overview (Circos Display):** Utilizing the R circlize package^28^, this module generates a polar circos plot providing a panoramic view of PTM distribution density, regional protein characteristics, and inter-sample differential regulation. It supports simultaneous comparison of multiple groups.

PTMExplorer incorporates the PTM-Disease Nexus module, which systematically aggregates data from three major disease-associated modification databases: PTMD^15^, qPTM^14^, and PhosCancer^13^. This allows users to perform co-localization analysis between their experimental differential PTMs and reported disease-related modification sites within the same interface, facilitating the assessment of candidate site clinical relevance. The platform supports eight model organisms (human, rat, mouse, fruit fly, worm, *E. coli*, yeast, and *Arabidopsis*). It accepts user-uploaded differential analysis tables, is compatible with output formats from mainstream search engines, and generates publication-ready figures.

## 2. Materials and Methods

### 2.1 Reference Feature Datasets

Reference feature data obtained directly by PTMExplorer from public databases comprises four parts (Table S1):

1. **UniProt Feature Annotations:** Sourced from the UniProt database (http://www.uniprot.org/), these cover fundamental protein sequence annotations, including Domains & sites, Molecule processing, PTM sites, Sequence information, Structural features, and Amino acid variations. These features intuitively display protein domain architecture, existing PTM annotations, and amino acid variation information.
2. **Sequence Databases:** UniProt reference proteome sequence databases for the eight commonly supported species.
3. **Protein 3D Structure Information:** 3D structural data originates from UniProt’s PDB cross-references, including PDB IDs and chain start/end positions. After retrieving structure files from the PDB database using the PDB ID, these files serve as the basis for subsequent 3D rendering.
4. **Modification-Disease Association Annotations (PTM-Disease Nexus):** Modification-disease association data from three major disease-related databases—PTMD, qPTM, and PhosCancer—have been uniformly curated and annotated. PTMD annotations retain their original PDA classification codes (e.g., a site marked U4D3P1A1C2N3 indicates counts for Up/Down/Presence/Absence/Create/Disrupt evidence types). For qPTM and PhosCancer, significance was determined using thresholds of p < 0.05 and Fold Change > 1.5 or < 0.67, labeling sites as Up (U), Down (D), or Not significant (N). Results display a summary across all curated datasets (e.g., U3D2E1 indicates upregulation in 3 datasets, downregulation in 2, and no significant change in 1).

### 2.2 Disorder and Surface Accessibility Calculations

Intrinsically disordered regions (IDRs) and surface accessibility are key factors influencing PTM occurrence probability.

IDR prediction is performed on the input amino acid sequence using flDPnn^26^, measured by the IDR score. A higher IDR score indicates greater disorder propensity, which correlates with a higher likelihood of PTM occurrence^7^.

Surface accessibility calculations are based on protein structure annotations downloaded from AlphaFold (https://alphafold.ebi.ac.uk/)^29^. The Relative Surface Accessibility (RSA) for each amino acid is calculated using FreeSASA^30^. Higher RSA values indicate greater exposure to the solvent surface, correlating with a higher likelihood of PTM occurrence^31^.

### 2.3 Experimental Dataset

To demonstrate the practical application of PTMExplorer, we utilized a previously published multi-modification dataset from hepatocellular carcinoma (HCC)^6^ (18 patients, 4D-Label Free, 9 modification types). Differential statistical testing was performed using t-tests between groups defined by tumor/cancer-adjacent status and HBV+/HBV-status. Significance was determined using thresholds of FC 1.5 and p-value 0.05. Two representative case study proteins were selected:

- **Q96ST2 (IWS1, 819 aa):** Used to demonstrate multi-group/temporal quantitative comparison and modification-protein joint analysis. Multiple sites were significantly upregulated in tumors, with most sites also having disease-related information recorded in PTMD, PhosCancer, and qPTM.
- **P31327 (CPS1, 1500 aa):** Used to demonstrate differential analysis across multiple modification types. This protein harbored detected sites for 9 modification types. While most sites were generally downregulated in tumor versus adjacent tissue, several succinylation and ubiquitination sites were significantly upregulated in HBV+ versus HBV-samples.

### 2.4 Data Input

User-input experimental data must be organized according to a specified format (see Supplementary Table S2), including: Protein Accession, Amino Acid Sequence, Modification Position, Modification Type (Category), Sample Quantitative Values, Differential FC, p-value, FDR, and Significance Determination. Additionally, a sample grouping file (containing Sample and Group columns defining sample relationships) is required.

PTMExplorer supports batch upload of multiple proteins (select species, upload experiment results and grouping file). It also supports a single-protein mode where users can either upload a FASTA file to define the reference sequence or use a protein ID for automatic sequence retrieval via ID mapping. If a single-protein FASTA file is uploaded, PTMExplorer performs a BLAST search against the selected species’ sequence database to precisely match feature data to the user-provided sequence.

After data upload, PTMExplorer automatically integrates all information, establishing sequence-feature correspondence. Quantitative values can be displayed as raw values or after Z-score normalization. In differential analysis results, FC is log2-transformed, p-value is -log10 transformed, and values are displayed using a color scale (red for upregulation, green for downregulation, grey for non-significant).

### 2.5 Technical Implementation of Three Visualization Modules

#### PTM Inspector

Built on ProtVista (developed by the UniProt consortium)^18^, the protein sequence is arranged horizontally, with corresponding feature tracks plotted below by position. PTMExplorer extends ProtVista by: (a) Supporting custom data sources, defining a universal data format for feature, variation, and proteomics data, allowing customization of categories, colors, and display information; (b) Providing dynamic category filtering, allowing users to select which feature types to display; (c) Enabling feature clicking to highlight covered regions, displaying precise positions, descriptions, and evidence details.

#### PTM 3D Locator

Built on the Nightingale nightingale-structure component^27^, using Mol* as the 3D rendering engine. The system retrieves structure files via PDB ID, then utilizes the PDBe-SIFTS service for precise UniProt-PDB residue mapping to correlate sequence positions with 3D spatial coordinates. The final output is an interactive 3D structure rendered in the web browser, allowing users to rotate, zoom, and highlight specific sites.

#### PTM Overview

Constructed using the R circlize package^28^ to generate polar circos plots. The entire protein sequence occupies a single sector, with the x-axis mapping amino acid positions. Each information type corresponds to one or more concentric tracks, drawn from outer to inner layers. Data processing involves two layers: (a) Data Layer: Uniformly parses various features into standard semantic types (domain/secondary structure/variation/modification/continuous value, etc.); (b) Rendering Layer: Maps each information type to an independent track, supporting empty track strategies, configurable order and height. An interactive annotation system manages labels and legends in partitioned areas for automatic alignment.

### 2.6 Other Calculations and Plotting

Demo data sourced from the literature retained its pre-processing and differential calculation results. However, re-verification of some data used the differential analysis module of BioLadder^25^ for calculating FC and p-values, and its boxplot module for plotting.

## 3. Results

### 3.1 Overall Design Framework of PTMExplorer

The core design philosophy of PTMExplorer is to integrate user-uploaded PTM experimental data (including sample/group quantitative values, differential FC, p-value, and significance calls) with UniProt sequence feature annotations, real-time computed sequence features, and disease-associated modification differential data. This integrated information is then simultaneously presented through three complementary visualization perspectives, enabling users to intuitively observe the positioning of differential modification sites within the protein sequence context, 3D structure, and panoramic distribution, thereby facilitating rapid assessment of their potential functional impact (Figure 1).

**Figure 1.**
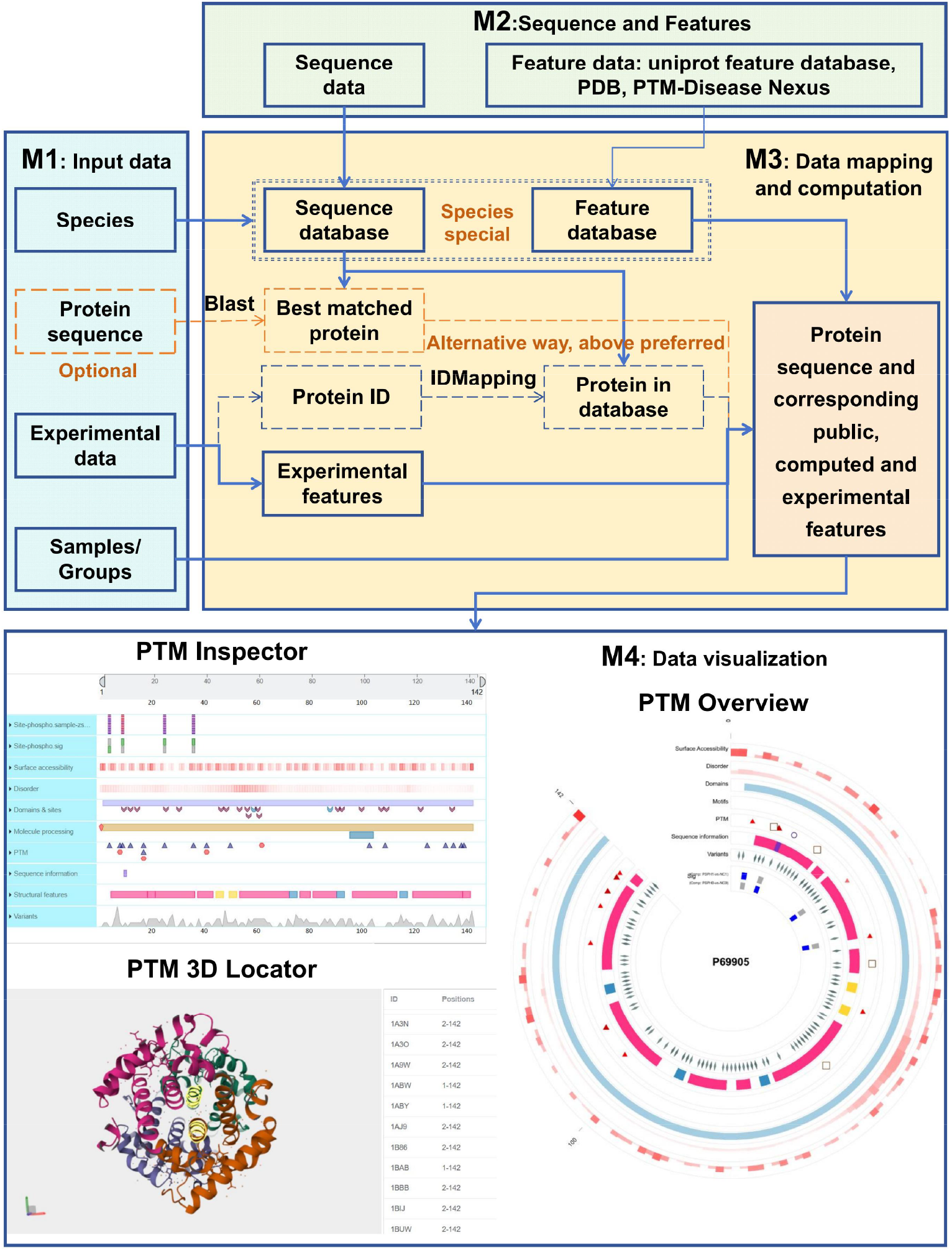
Schematic overview of the PTMExplorer workflow. Divided into four modules. M1: Input data, comprising user-provided information including experimental data, sample/group information, species selection, and optional protein sequence input for single-protein mode. M2: Sequence and Features, encompassing public feature data including species-specific sequence databases, feature annotations, PDB structure data, and the PTM-Disease Nexus. M3: Data matching and calculation, involving matching input experimental data (protein ID or sequence) against feature databases and calculating surface accessibility and disorder metrics. M4: Data visualization, presenting three distinct visualization methods.

### 3.2 Quantitative Comparison for Multi-Group/Temporal Data

PTMExplorer can display sample or group quantitative data alongside differential analysis results. Using the HCC multi-modification study data (Table S2), the IWS1 protein (Q96ST2) serves as a case study. IWS1 is a crucial transcriptional regulator, playing a key role in the composition of the RNA polymerase II (RNAPII) elongation complex and the production of mature mRNA transcripts^32^. The study identified 23 phosphorylation sites on this protein, many of which were significantly upregulated in cancer samples compared to adjacent normal tissue (Fig 2A). We focus here on eight high-confidence upregulated sites (Fig 2B). All eight sites are located in regions of high surface accessibility, and the first six exhibit high disorder scores (Fig 2A, 2B). Within the PhosCancer database, most of these sites are reported as upregulated across various cancer types; in qPTM, both up- and down-regulation are observed; six sites are recorded in PTMD, labeled ‘N’ (Disruption), indicating that mutation events at these sites potentially disrupt one or more PTM sites or reduce protein PTM levels in disease^15^.

**Figure 2.**
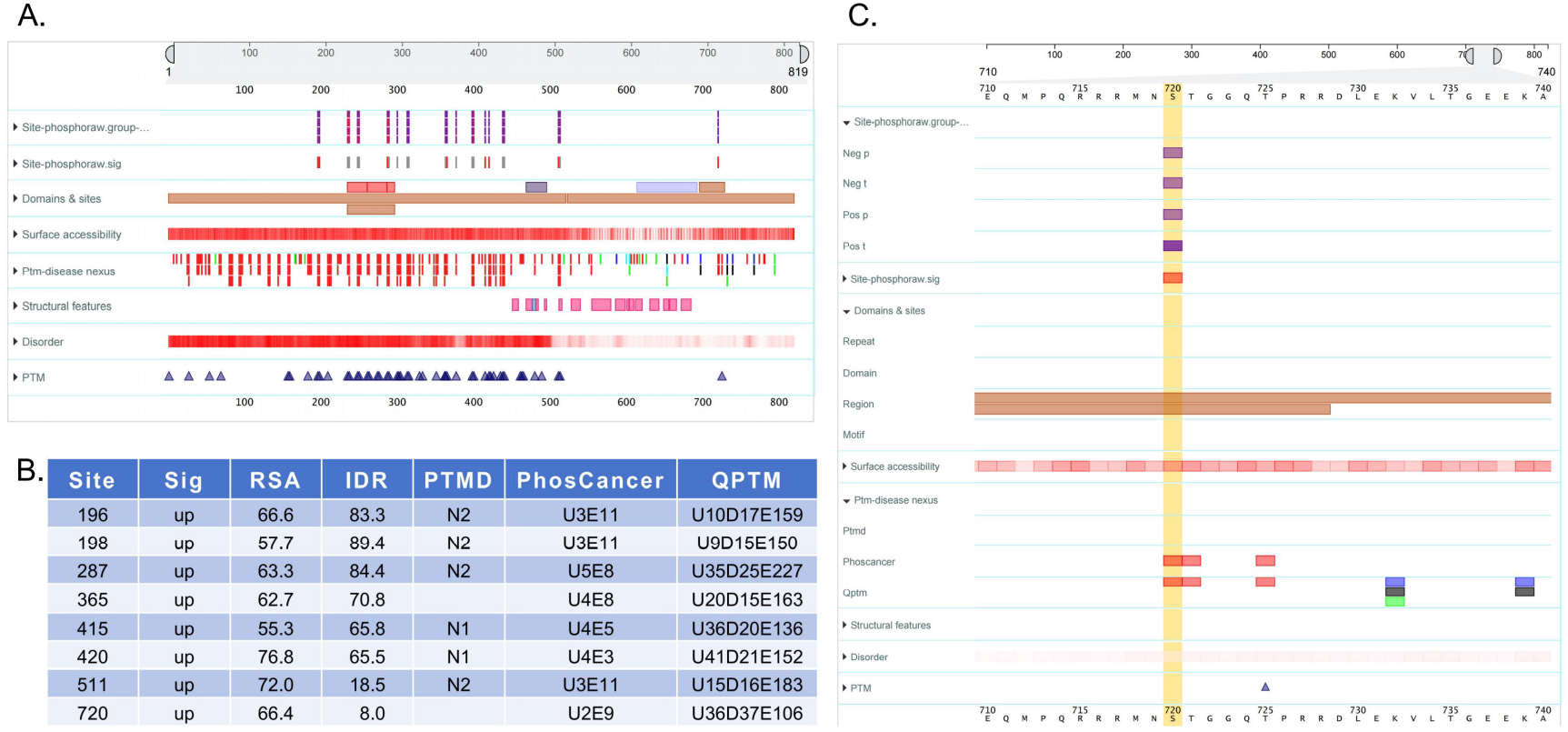
Multi-group quantitative comparison for protein IWS1 (Q96ST2). (A) PTM Inspector full-protein view. The leftmost two rows show grouped phosphorylation site quantitative values (without Z-score transformation) and the significance of cancer versus adjacent normal tissue differences. Lower tracks display reference feature data. (B) Detailed data for eight high-confidence upregulated sites. RSA indicates surface accessibility; IDR indicates disorder score. The last three columns show annotations from the three integrated PTM-Disease Nexus datasets. (C) Detailed information for the Ser720 phosphorylation site. This is an Akt1/Akt3 phosphorylation site. Its ABS is significantly upregulated in cancer versus adjacent tissue (FC = 1.61, p = 0.041). Its IDR score of 8.0 places it in a low-disorder region, suggesting high RSA alone can drive phosphorylation. PhosCancer marks it as U2E9. Detailed information reveals upregulation reported in UCEC, OV, LUAD, LUAD_cnHPHP, LSCC datasets, but non-significance in HCC and HCC_cnHPHP.

Notably, phosphorylation at Ser720 is significantly upregulated in cancer tissue compared to adjacent normal tissue (FC = 1.61, p = 0.041). Its IDR score is only 8.0, placing it in a low-disorder region—a stark contrast to the first six high-IDR upregulated sites. This suggests that high surface accessibility (RSA) alone can drive phosphorylation, and disorder is not a prerequisite. This site is a known Akt1/Akt3 phosphorylation target. Literature reports that in non-small cell lung cancer,

Akt-mediated phosphorylation of IWS1 recruits SetD2 to increase H3K36me3 modification, further recruiting the MRG15/PTB complex to regulate alternative splicing of FGFR2 (IIIb→IIIc), promoting tumor progression^32,33^. PhosCancer marks this site as U2E9. Linking to the PTM-Disease Nexus reveals detailed information: it is reported as upregulated in datasets for GBM, BRCA, etc., but non-significant in HCC and other cancers (Fig 2C).

These results demonstrate that PTMExplorer provides users with convenient multi-dimensional feature analysis for modification data, effectively helping researchers anchor experimental differential sites to reported disease-associated sites. Combined with features like IDR and RSA, it facilitates rapid assessment of site functional potential and prioritization of clinically relevant candidates.

### 3.3 Differential Analysis Across Multiple Modification Types

PTMExplorer supports joint differential analysis across multiple modification types. Taking the P31327 protein (gene name CPS1, Carbamoyl-phosphate synthase [ammonia], mitochondrial) as an example (Table S3), the HCC dataset identified 9 modification types across 61 modification sites, totaling 198 modifications (see Supplementary Table S3). Notably, lysine 55 (K55) was found to harbor six different modifications (Acetylation, Crotonylation, Lactylation, Succinylation, Malonylation, β-hydroxybutyrylation), and several other sites also exhibited multiple modifications, suggesting potential crosstalk.

Overall comparison between tumor and adjacent normal tissue revealed 123 downregulated modification sites (62% of total) and no upregulated sites, consistent with the approximately 2.8-fold downregulation of the CPS1 protein itself in tumors (FC = 0.35, p < 0.01), likely reflecting a protein abundance effect (Fig 3A). However, in the comparison between HBV-positive and HBV-negative samples, seven modification sites were upregulated (three succinylation sites, three ubiquitination sites, and one crotonylation site, Fig 3A). Figure 3B uses a circos plot to provide an overview of succinylation differences in tumor and HBV comparisons.

**Fig. 3.**
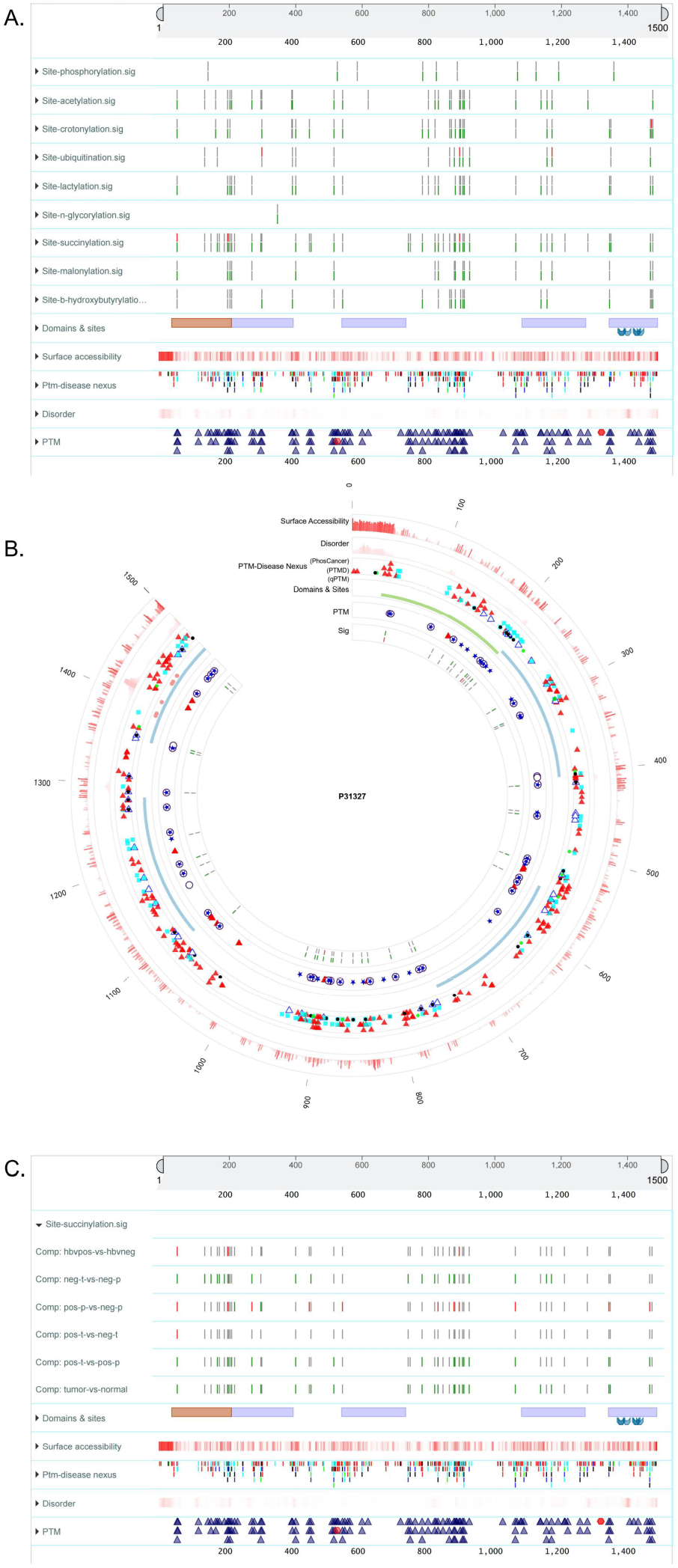
Multi-modification differential analysis of protein P31327. (A) PTM Inspector view showing the differential significance of nine modification types across comparison groups. Succinylation (succi) was predominantly downregulated in tumor versus adjacent normal (green), whereas multiple succinylation sites were upregulated in HBV^+^versus HBV^−^ (red). (B) PTM Overview circos plot providing a panoramic view of succinylation distribution density and differential significance across protein regions. Succinylation sites were concentrated in the N-terminal and central regions of the protein. (C) PTM Inspector displaying the differential analysis results of succinylation across six comparison pairs. The direction and magnitude of succinylation changes were clearly dependent on both tissue type (adjacent normal versus tumor) and HBV infection status, highlighting the context-specific nature of this modification.

Strikingly, in adjacent normal liver tissue, the HBV-positive group showed higher succinylation levels at 11 CPS1 sites compared to the HBV-negative group (Pos_P vs Neg_P, Fig 3C). However, in tumor tissue, only one succinylation site remained upregulated between HBV-positive and negative groups, with others showing no significant difference (Pos_T vs Neg_T, Fig 3C). This observation suggests that the regulatory effect of HBV infection on CPS1 succinylation exists in normal liver tissue but may be masked during tumorigenesis by stronger metabolic reprogramming effects—a hypothesis requiring validation in larger, independent cohorts.

These results illustrate that in multi-modification joint analysis, PTMExplorer can rapidly present the global modification landscape of a protein, revealing its differential regulation patterns under distinct biological conditions.

### 3.4 Modification–Protein Joint Analysis: Raw vs. Protein-Normalized Modification Quantification

PTMExplorer also supports joint analysis of modification and protein abundance. To distinguish the two quantification layers, we define raw modification abundance as the directly measured MS intensity of a modified site, and protein-normalized modification abundance (hereafter normalized abundance) as the raw abundance divided by the abundance of the corresponding protein. Raw abundance reflects the absolute level of a modification in the sample and is directly tied to its functional readout. Normalized abundance, by contrast, isolates the regulatory contribution of the modification machinery by removing the confounding effect of protein expression changes.

Comparing these two quantities together with protein abundance can reveal distinct regulatory modes. When protein and normalized abundance change in the same direction, the modification change is reinforced by both protein-level and modification-level regulation. When they change in opposite directions, modification-level regulation counteracts the protein change, and the net direction of raw abundance is determined by whichever influence dominates.

We illustrate this framework using IWS1 (Q96ST2) (Table S4). The protein showed a mild, statistically significant decrease in tumor versus adjacent tissue (FC = 0.68, p < 0.01, though FC did not pass the threshold). Strikingly, among the 23 identified phosphorylation sites, 21 showed increased normalized abundance in tumors, of which 8 also had significantly increased raw abundance (modification-regulation-driven), while the remaining 13 showed no significant change in raw abundance (protein-abundance-driven) (Fig 4A). This pattern indicates that despite a slight reduction in total IWS1 protein, its phosphorylation was under strong active regulation, with eight sites becoming more abundant in absolute terms due to increased modification density.

**Fig. 4.**
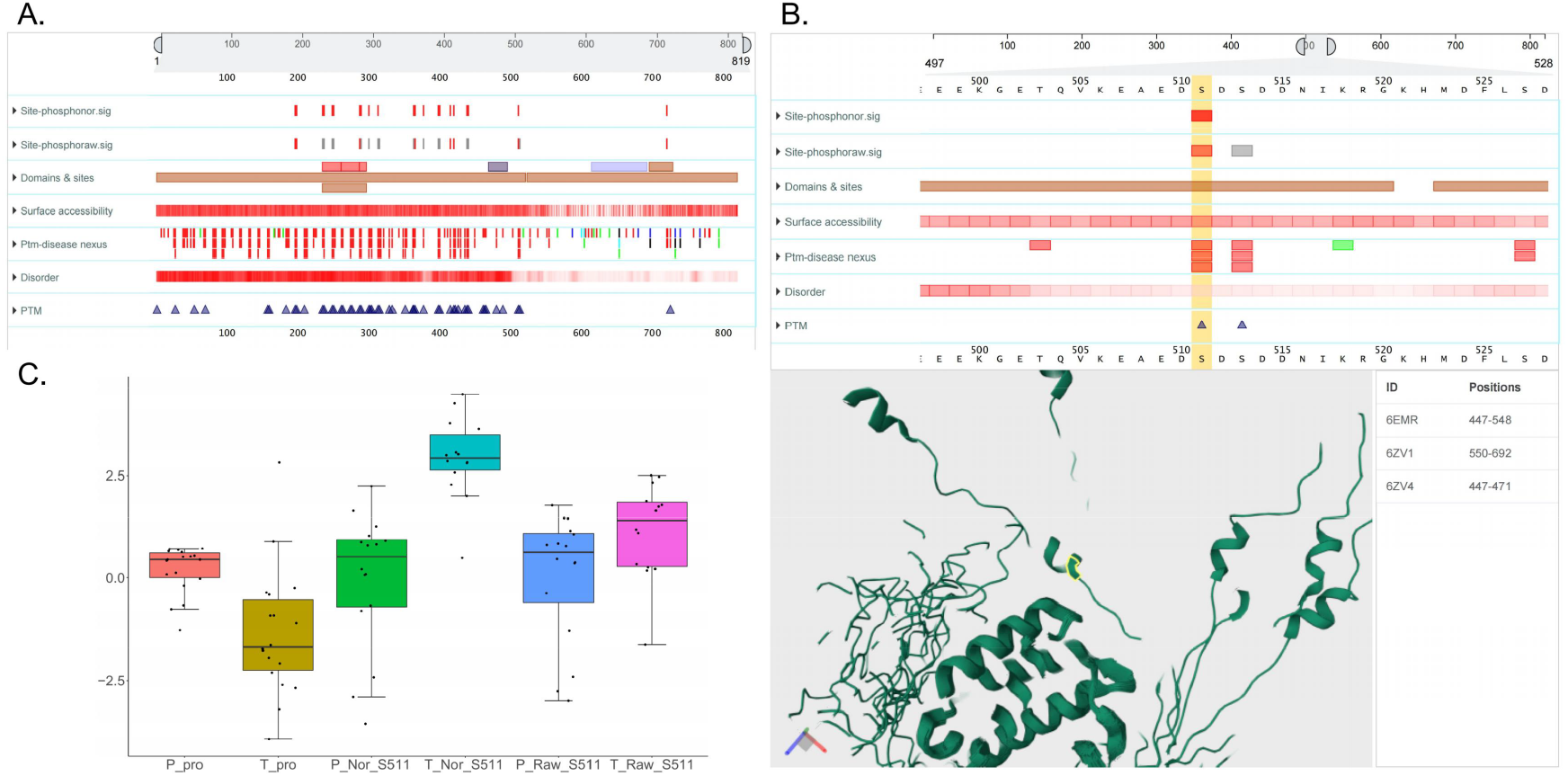
Integrated protein abundance and PTM-site analysis of protein Q96ST2 (IWS1). (A) Comparison of raw and protein-normalized modification abundance across 23 phosphorylation sites. Among them, 21 sites showed significantly increased normalized abundance in tumors (red), of which 8 also exhibited significantly increased raw abundance, while the remaining 13 showed no significant change in raw abundance (gray), reflecting the independent contribution of post-translational regulation. (B) Significance overview of the Ser511 phosphorylation site on IWS1 along with its associated reference features. The PTM 3D Locator panel below provides the spatial localization of this site within the three-dimensional structure. (C) Paired boxplot comparison of protein abundance, normalized modification abundance, and raw modification abundance for Ser511. P_pro, protein abundance in adjacent normal tissue; T_pro, protein abundance in tumor tissue; P_norm_S511 / T_norm_S511, protein-normalized modification abundance of Ser511; P_raw_S511 / T_raw_S511, raw modification abundance of Ser511. Protein abundance was mildly decreased in tumors (FC = 0.68, p < 0.01, FC below threshold); normalized abundance was strongly upregulated (FC = 6.04, p < 0.01); raw abundance was also significantly elevated (FC = 1.86, p < 0.05), indicating that despite a modest reduction in total IWS1 protein, phosphorylation at Ser511 was actively enhanced through post-translational regulation.

Figure 4B details Ser511 as an example. This site resides in a beta-sheet region and exhibits high surface accessibility. Boxplots of protein abundance, normalized abundance, and raw abundance for Ser511 are shown in Figure 4C. Protein abundance was mildly lower in tumors (FC = 0.68, p < 0.01); normalized abundance was strongly upregulated (FC = 6.04, p < 0.01); and raw abundance was also significantly elevated (FC = 1.86, p < 0.05). This demonstrates that even when the protein declines, the modification density at Ser511 is actively enhanced, resulting in a net increase in the absolute phosphorylated population.

These results highlight that PTMExplorer’s dual-layer quantification view can reveal cases where post-translational regulation compensates for or overrides protein-level changes, providing a more complete picture of signaling activity than either layer alone.

## 4. Discussion

### 4.1 Integration of User-Defined PTM Data with Multi-Dimensional Protein Contexts

PTMExplorer enables the simultaneous projection of user-defined differential PTM data onto multi-dimensional protein contexts—including domain architecture, functional sites, three-dimensional spatial location, disorder propensity, surface accessibility, and disease associations—within a single online interactive interface. This design directly addresses a common challenge in modification proteomics: prioritizing functionally important candidate sites from a large list of differentially modified ones.

Compared to existing tools, PTMExplorer offers several distinctive features. First, it systematically integrates UniProt functional annotations, IDR prediction (flDPnn), surface accessibility (FreeSASA/AlphaFold), and PDB structural information within the same sequence tracks, moving beyond the single-annotation-track display typical of most tools. Second, the three complementary visualization perspectives work together: PTM Inspector reveals site-level sequence context and functional annotations, PTM 3D Locator provides spatial localization, and PTM Overview delivers a panoramic view of regulatory patterns across modification types and group comparisons. Third, the platform accommodates multiple experimental scenarios, including multi-modification analysis, multi-group comparisons, temporal series, and joint proteome and post-translational modification analysis. The dual-layer display of raw and protein-normalized modification abundance helps disentangle modification-level regulation from protein abundance effects.

### 4.2 Methodological Considerations and Current Scope

Several points should be considered when interpreting results generated by PTMExplorer.

#### Three-dimensional structure representation

The 3D structures displayed in PTMExplorer are obtained from the PDB or AlphaFold and represent the canonical form of the protein. Although some PDB entries do contain modified residues, the specific modification sites identified in a given experiment are generally absent from the available structural models. As a result, the spatial context provided should be understood as the framework of the unmodified backbone, rather than a precise representation of the post-translationally modified conformation. Upon modification at a particular site—such as the introduction of a negatively charged phosphate group or the removal of a positive charge by acetylation—local or even global conformational changes may occur. Predicting such post-modulation structural rearrangements remains an open challenge across the field, as even state-of-the-art methods like AlphaFold3 produce static folding snapshots rather than dynamic ensembles. Future integration of molecular dynamics simulations or machine-learning-based conformational sampling may help bridge this gap as the relevant methods mature.

#### Detection-level limitations in multi-modification co-occurrence

Current bottom-up proteomics workflows rely on enzymatic digestion followed by peptide-level identification. This approach cannot determine whether multiple modification sites on the same protein coexist on a single protein molecule unless the sites reside on the same identified peptide. Users should bear this limitation in mind when interpreting potential crosstalk between modification sites.

#### Coverage of disease-associated modification databases

Although the PTM-Disease Nexus integrates three databases—PTMD, qPTM, and PhosCancer—over 400 types of PTMs are known to exist in nature, and systematic functional annotations remain scarce for many species- and disease-specific modifications. As the field continues to accumulate modification proteomics data, PTMExplorer will incorporate additional functional and disease-associated annotations.

### 4.3 Applications

We have outlined three main application scenarios for this platform: comparative display of quantitative data (including multi-group and temporal comparisons), visualization of differential analysis across multiple modification types, and joint PTM–proteome analysis. These three categories cover the majority of current modification-oriented studies. Future research paradigms involving modifications can also adopt this platform for visualization after appropriate data formatting.

The observations described above are derived from the specific dataset visualized through PTMExplorer and represent inferences consistent with the data as presented. Biological systems are inherently complex, and the relationships suggested here may involve additional layers of regulation not captured in this analysis. These findings should therefore be regarded as hypotheses warranting further experimental validation or complementary investigation.

Beyond modification-specific applications, the design architecture of PTMExplorer is readily extensible. By adjusting the Category field in the input file, the visualization target can be switched from modification sites to amino acid mutations or identified peptides. For example, setting Category to mutation types will display mutation positions within the protein sequence and 3D structure along with their potential functional impact. Replacing the single amino acid in the Sequence column with a peptide sequence (with the display field automatically switching from Site to Peptide) allows visualization of identified peptide distributions from peptide-centric experiments such as Lip-MS. This flexibility makes PTMExplorer adaptable to a wide range of applications in proteomics and modificomics.

### 4.4 Platform and Accessibility

PTMExplorer is integrated into the BioLadder bioinformatics cloud platform^25^. No local software installation is required; it is accessible via a web browser at https://www.bioladder.cn/PTMExplorer/.

## 5. Conclusion

In this study, we developed PTMExplorer—an interactive visualization platform that systematically integrates proteomic PTM differential data with multi-dimensional protein functional and structural features (https://www.bioladder.cn/PTMExplorer/). Through three complementary modules—PTM Inspector, PTM 3D Locator, and PTM Overview—PTMExplorer simultaneously presents modification data from one-dimensional sequence, three-dimensional space, and panoramic circos perspectives. It addresses four core deficiencies in existing tools: lack of multi-dimensional feature integration, absence of 3D structural localization, inability to import user differential data, and missing disease-site co-localization. Using a multi-omics hepatocellular carcinoma dataset as a case study, we demonstrated PTMExplorer’s practical value in screening key biomarkers, revealing multi-modification coordination mechanisms, and distinguishing patterns between absolute and relative quantification.

## Acknowledgments

This work was supported by National Key R&D Program: Key Technologies for Fine-Grained Analysis and Secure Sharing of High Spatiotemporal-Throughput Medical Data (2024YFE0202700, 2024.12-2027.11), and the National Natural Science Foundation of China (Grant No. 82472946).

## Declarations

All authors have read and approved the final version of this manuscript. This work has not been published or accepted for publication elsewhere.

